# Kindlin Assists Talin to Promote Integrin Activation

**DOI:** 10.1101/662163

**Authors:** Z. Haydari, H. Shams, Z. Jahed, M.R.K. Mofrad

## Abstract

Integrin αIIbβ3 is a predominant type of integrin abundantly expressed on the surface of platelets and its activation regulates the process of thrombosis. Talin and kindlin are cytoplasmic proteins that bind to integrin and modulate its affinity for extracellular ligands. While the molecular details of talin-mediated integrin activation are known, the mechanism of kindlin involvement in this process remains elusive. Here, we demonstrate that the interplay between talin and kindlin promotes integrin activation. Our all-atomic molecular dynamics simulations on complete transmembrane and cytoplasmic domains of integrin αIIbβ3, talin1 F2/F3 subdomains, and kindlin2 FERM domain in an explicit lipid-water environment over microsecond timescale, unraveled the role of kindlin as an enhancer of the talin interaction with the membrane proximal region of β–integrin. The cooperation of kindlin with talin results in a complete disruption of salt bridges between R995 on αIIb and D723/E726 on β3. Furthermore, kindlin modifies the molecular mechanisms of inside-out activation by decreasing the crossing angle between transmembrane helices of integrin αIIb-β3, which eventually results in parallelization of integrin dimer. In addition, our control simulation featuring integrin in complex with kindlin reveals that kindlin binding is not sufficient for unclasping the inner membrane and outer membrane interactions of integrin dimer, thus ruling out the possibility of solitary action of kindlin in integrin activation.

**Statement of Significance:** Using the newly solved crystal structure of kindlin, we investigated, for the first time, the molecular mechanism of kindlin-mediated integrin activation through simultaneous binding of talin and kindlin. We demonstrate in atomist details how kindlin cooperates with talin to promote the activation of integrin αIIb-β3.

## Introduction

Integrin plays a central role in regulating cell-matrix and cell-cell adhesion and is crucial for various signaling pathways involved in cell migration, proliferation, and differentiation (1,2). Integrins are heterodimeric proteins composed of α and β subunits that associate noncovalently. Each subunit consists of a large extracellular ectodomain (ECD), a single pass transmembrane helix (TM), and a short cytoplasmic tail (CT) (Fig. 1A) (3,4). There are 24 different combinations of integrin α and β subunits. Integrin αIIbβ3 is one of the well-known types that is only expressed in platelets and is necessary for the hemostatic function of platelets (5,6). Integrins mediate bidirectional, inside-out and outside-in signaling across the membrane. Inside-out signaling involves the interaction of intracellular proteins with the integrin CT, which modulates integrin’s affinity for extracellular ligands (7,8).

**Figure1:**
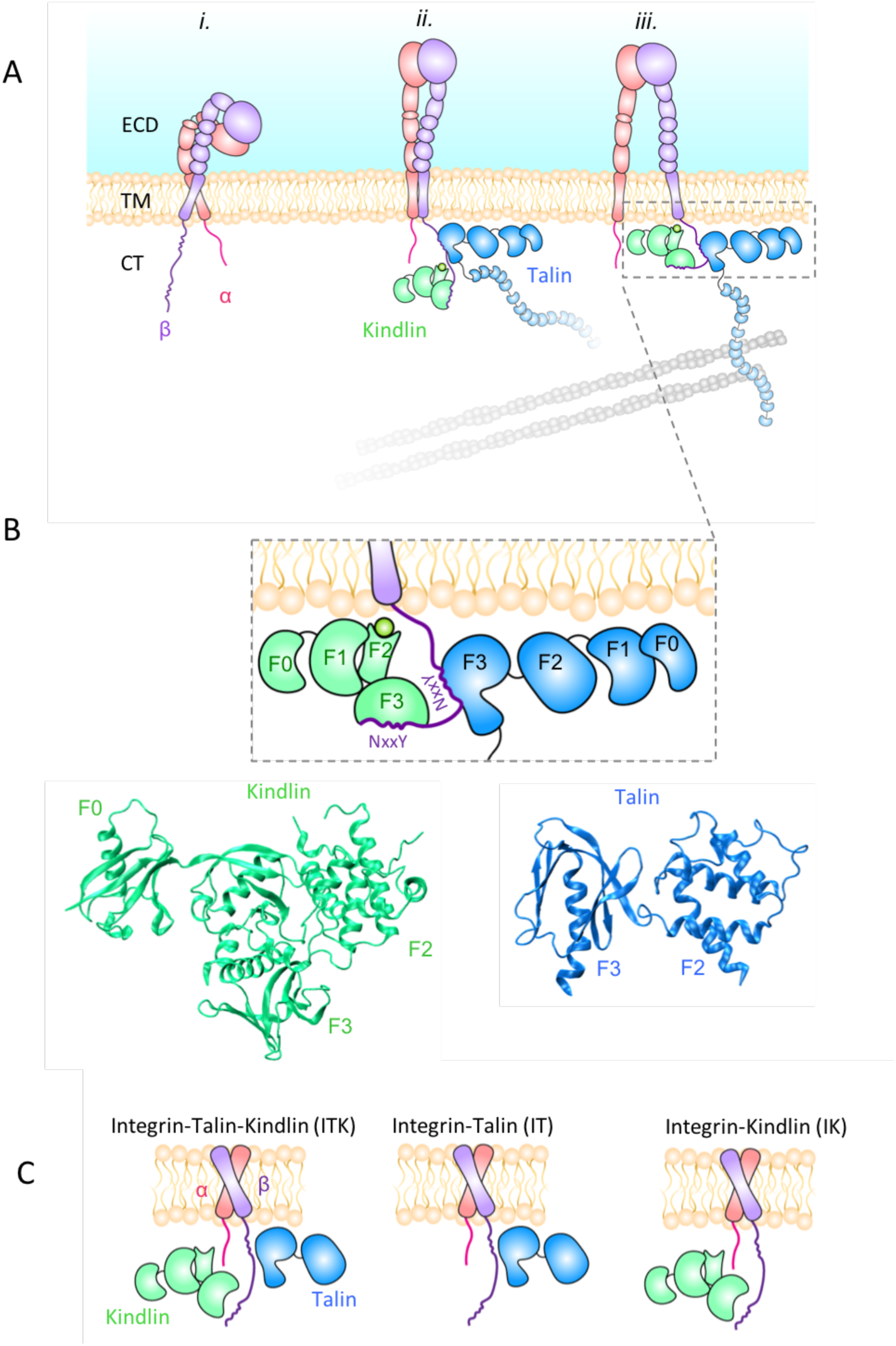
Cytoplasmic interaction with the integrin tail mediate inside-out signaling. **(A)** Schematic representations of the three conformational states of integrin: i) bent-closed (inactive) ii) extended-closed iii) extended-open (activated). Talin and kindlin are both involved in integrin activation by binding to the cytoplasmic domain of β. Talin association to the actin cytoskeleton provides a linkage between the extracellular environment and the cytoskeleton. **(B)** The F3 subdomain of talin and kindlin bind to the membrane proximal and membrane distal regions of β–integrin, respectively. (top). The crystal structure of kindlin2 (F0-F3 subdomains) and talin (F2-F3 domains). **(C)** Schematic representations of the three molecular models used in our simulations: integrin in complex with talin and kindlin (ITK), integrin in complex with talin (IT), and integrin in complex with kindlin (IK).

Three distinct conformational states of integrin have been determined: bent-closed or the resting state, extended-closed, and extended-open (Fig. 1A) (6). Integrin activation entails a conformational transition of integrin from a low-affinity (bent-closed) to a high affinity state (extended-open) (Fig. 1A) (9,10). It was previously believed that talin was both necessary and sufficient for inducing integrin activation (11,12), whereas recent studies contradict this notion (13–15) and suggest that kindlin is also needed as a co-activator of integrin (13,16,17). For example, Moser et al. showed that kindlin3 knockout cells results in inactivated integrins despite presence of talin (18). Moreover, kindlin-associated diseases like Leukocyte Adhesion Disease III (LADIII), reflecting deficiencies of integrin function, show the necessity of kindlin in integrin activation (5).

Both talin (talin-1 and −2) and kindlin (kindlin-1, −2, and −3) isoforms include a FERM (4.1 protein, ezrin, radixin, moesin) domain, which is composed of four structural subdomains F0-F3 (19). Kindlin also features a pleckstrin homology (PH) domain inserted into its F2 subdomain, which has been shown to directly interact with the lipid membrane (Fig. 1B) (20). Talin and kindlin both bind to the CT domain of integrin through their F3 subdomains. Specifically, talin binds to the NxxY motif at the membrane proximal region, while kindlin associates with this motif at the membrane distal part of CT suggesting that kindlin and talin can simultaneously bind to integrin tail and trigger activation (21). The molecular mechanisms of the interplay among talin and kindlin for inducing integrin activation are not yet well understood.

Recent experimental and computational studies have revealed important molecular details of integrin inside-out activation. These studies have suggested that the inner membrane clasp (IMC) between the CT domains of integrin subunits is critical for maintaining the closed state of integrin and unclasping induced by talin or other activators triggers activation (22–24), and that opening of IMC results in an increase in the crossing angle between α and β monomers (25–27). However, all computational studies on integrin-mediated inside-out signaling to date have overlooked the role of kindlin due largely to lack of sufficient structural information (7,28–30). In the present study we use the recently solved crystal structure of kindlin (31) to investigate the molecular mechanism of integrin activation through simultaneous binding of talin and kindlin. We use all-atomic microsecond-scale molecular dynamic simulations of αllbβ3 TM/CT structure in an explicit lipid-water environment under the following three conditions: in complex with talin1 F2-F3 subdomains (IT), with kindlin2 (IK), and with both (ITK) to uncover the role of kindlin in integrin activation (Fig. 1C). Specifically, we explore how talin and kindlin binding affect allosteric transition of forces across the integrin molecule, i.e. from CT to TM, and eventually to the ectodomain. Our results show that kindlin2 cooperates with talin1 to facilitate integrin αllbβ3 activation by enhancing talin1 interaction with the membrane proximal region of β3-integrin.

## Materials and Methods

### System Setups

A system consisting of TM/CT domains of integrin αIIbβ3 (PDB: 2KNC (32)) embedded in the plasma membrane, talin1 F2/F3 subdomains (PDB: 3IVF (20)), and kindlin2 FERM domain (PDB: 5XQ0 (33)) (ITK) was built using VMD. Two control systems in which integrin αIIbβ3 was simulated in complex with either talin1 F2/F3 subdomains (IT) or kindlin2 FERM domain (IK) were also built. Two regions of the kindlin2 FERM domain are missing in solved crystal structure: residues 168–217 in the F1 subdomain and residues 337–512 in the F2 subdomain. The three systems were solvated in explicit water using NMAD solvated package. The TIP3 water model was used for solvation (34). However, the water molecules inside the lipid membrane were removed. Then the systems were neutralized and ionized with 150 mM of KCl. The number of atoms reached 744946 in ITK, 556876 in IT, and 625742 in IK.

### MD simulations

We performed all-atom molecular dynamics (MD) simulation to investigate the conformational changes of integrin αIIbβ3 upon simultaneous interaction with talin1 and kindlin2 (ITK) and compared them to the two control simulations (IT and IK) (Fig. 1A). Molecular dynamics simulations were carried out using NAMD and CHARMM36 force field (35). Molecular visualization and analysis were performed using Visual Molecular Dynamics (VMD) package (36). Periodic boundary conditions were used in all three dimensions and a 2 fs time step was used in all simulations. The Langevin piston Nose-Hoover pressure control algorithm, and the Langevin damping thermostat for temperature control were used (35). Pressure was maintained at 1 bar and temperature at 310 K with a damping coefficient of 5/ps.

Each trial was initially minimized for 100,000 timesteps using the conjugate gradient and line search algorithm to relax the structures and remove all bad contacts. Following the minimization process, each configuration was equilibrated for 5 ns or longer until equilibrium was reached. Fully equilibrated structures were then used in the final production simulations that ran for 1000 ns for ITK and IT, and 760 ns for IK.

### H-bond calculations

VMD hbonds plug-in (version 1.2) (36) were used to calculate the number of hydrogen bonds (hbonds) between regions of interest. The cutoff distance and angle were set to 4 A ° and 20, respectively. The density- and time-dependent plots were all prepared in R matrixStats and gplot (37,38).

### Principal component analysis (PCA)

The PCA analysis was used to examine time evolution of the integrin structure along all trajectories and understand the conformational differences induced by cytoplasmic interactions. The principal components were defined as the orthogonal axes of maximal variance and identified by superpositioning the structures on the invariant core and calculating the variance using the tools introduced by the Bio3D package (39).

### Residue cross correlation

To examine how the atomic fluctuations and displacements within the integrin heterodimer are correlated, we used the cross-correlation function in the Bio3D package (39). The matrix of all pairwise cross-correlations between residues were visualized using the dynamical cross-correlation map.

### Cross correlation function

The cross correlation function (CCF) in R was used to understand the relationship between two time series representing different features along the trajectories. The first argument was treated as the predictor (cause) or the second argument. The lag period indicates when the effect of a change in one feature is reflected in the other feature (39).

### Solvent accessible surface area (SASA)

The solvent-accessible surface area of each molecule was calculated using the measure SASA command in VMD (36). This command

### Force distribution analysis

To monitor changes in the internal forces of the integrin heterodimer upon cytoplasmic interactions, we performed Time-resolved Force Distribution Analysis (TRFDA) implemented in the GROMACS software package (40). Atomic pairwise forces were calculated for all residues of both integrin subunits and included only the non-bonded electrostatic interactions acting on each residue. The comparison between force propagation along the transmembrane domains of integrins indicated the allosteric signal propagation across the membrane. The punctual stress measured for alpha- and beta-subunits is the sum of absolute values of scalar pairwise forces exerted on each atom.

### Structural alignment

To determine the changes in the conformation of kindlin and talin upon interactions with integrin, we compared the initial structures of these molecules with the final structures after ∼1000ns of simulation. The final structures of talin in the ITK or IT simulations were aligned to the initial structure of talin using the *struct.aln* function in the bio3D package in R (39). Similarly, the final structures of kindlin in the ITK or IK simulations were aligned to the initial kindlin structure. To quantify the distances between residues in the two aligned structures, a difference vector was calculated between the two structures using the *dist.xyz*function in R (39).

## Results

To explore the role of kindlin in the activation of integrin as part of the integrin-mediated inside-out signaling, we developed all-atomic microsecond-scale molecular dynamic simulations of αllbβ3 using an explicit lipid-water environment under three distinct scenarios, namely integrin in complex with talin1 F2-F3 subdomains (IT), with kindlin2 (IK), and with both talin and kindlin (ITK).

### The interactions of talin1 and kindlin2 with the CT of integrin αIIbβ3 regulate its conformation

The mechanism of interaction of talin1 and kindlin2 with integrin β3 can be characterized based on the conformational transitions of the integrin heterodimer. These conformational changes were determined by comparing the first and last frames of our microsecond molecular dynamics simulations, featuring identical starting conformation of integrin for all scenarios (Fig. 2A). The crossing angle between the transmembrane regions of integrin monomers was notably increased in IT simulations changing the shape to open scissors (see below for further details). Unexpectedly, the final shape of integrin in the ITK simulation was changed to closed scissors (Fig. 2A). No significant change in the crossing angle was observed in the IK simulation.

**Figure 2:**
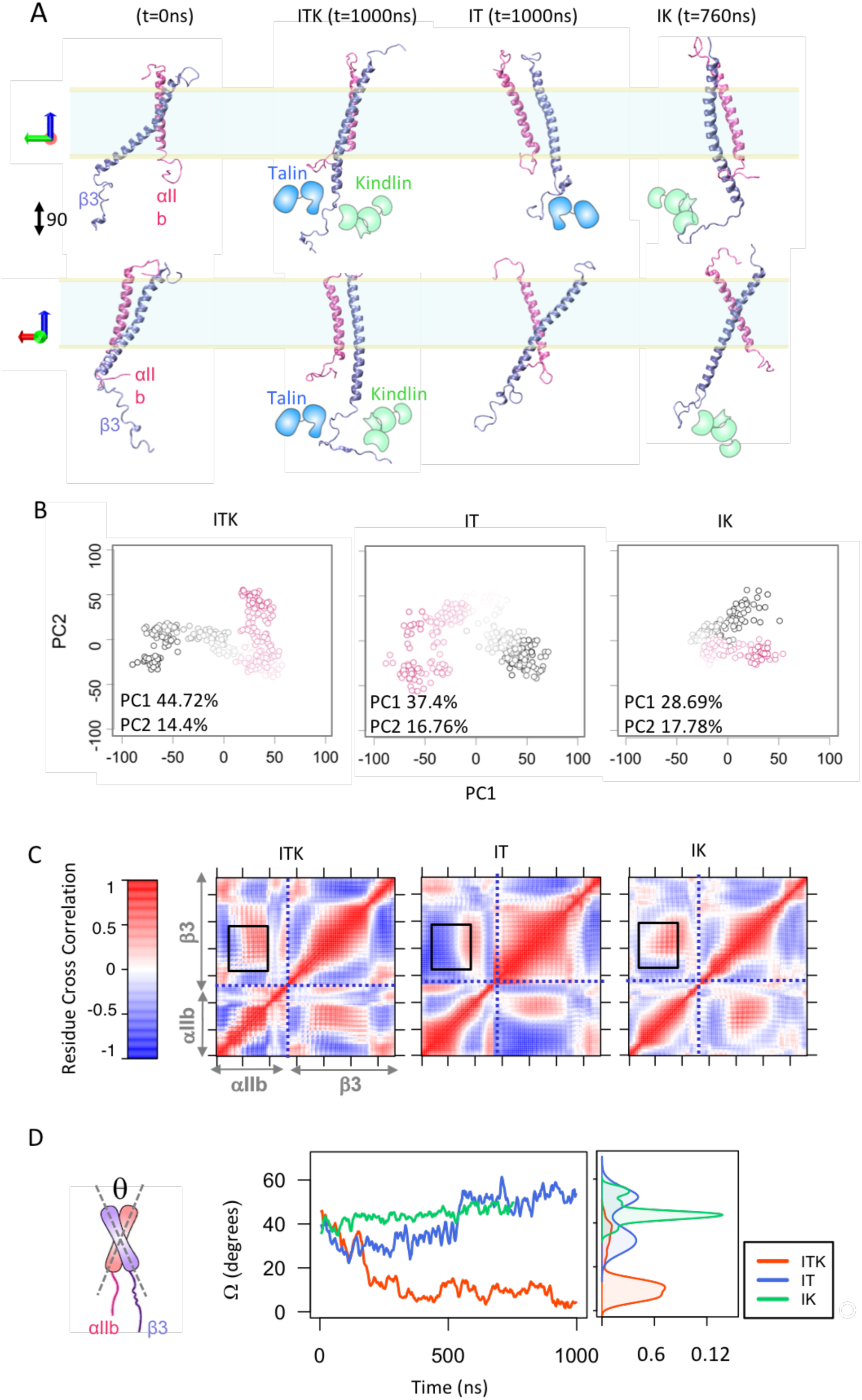
Conformational changes of integrin aIIbb3. **(A)** Schematics of talin and kindlin overlaid with the cartoon representations of the integrin αIIb (mauve) β3 (iceblue) heterodimer in two different views (90 degrees) (left top and bottom). Snapshots of the trajectories of the integrin αIIbβ3 dimer for the last simulation frame of the ITK (t=1000ns), IT (t=1000ns), and IK (t=760ns) simulations are shown. **(B)** The principal component analysis (PCA) of all trajectories. Each point is a structural state in the PC1-PC2 space. Points corresponding to the beginning of all trajectories are shown in red, middle frames are represented by smooth color change to white and end frames are illustrated in black. **(C)** Residue cross correlation heatmaps of the integrin αIIb and β3 helices averaged over simulation time. The αIIb and β3 regions are indicated on the heatmaps. Black boxes indicate the TM regions of integrin dimer. A representative heatmap is shown for each of the three ITK, IT, and IK simulations. **(D)** A schematic representation of the integrin dimer showing angle θ between the αIIb and β3 helices (right). The time plot and density plot of angle θ for the ITK, IT, and IK simulations are shown in red, blue, and green respectively.

Conformational transitions of the integrin dimer in response to cytoplasmic interactions with talin1 and/or kindlin2 were quantified via a principal component analysis (PCA). For PCA calculations, the invariant core of the structure in each simulation was identified, the distribution of structures was extracted from each trajectory, and conformational differences were quantified for equivalent residues. The first two principal components accounted for over 50% of the variance in ITK and IK, while in IT this number reduced to 40%. Since the contribution of all other principal components was relatively minimal, the structural distribution was projected in the PC1-PC2 space as shown in Fig. 2B. Three distinct clusters exist in IT and ITK, each representing structurally similar states, whereas only two are recognizable in IK. Moreover, final states of integrin in IT and ITK diverged significantly relative to the first frame.

Residue cross correlation analysis between all residue pairs of the integrin αIIb and β3 in ITK, IT, and IK was performed to examine how the fluctuations within the integrin dimer are correlated. Interestingly, we observed a coupling between the transmembrane regions of integrin αIIb and β3 (black boxes in Fig. 3C), which were dominantly correlated in ITK, but anti-correlated in IT. The level of residue correlations were notably lower in IK simulations resulting in a relatively lighter heat map (Fig. 3C). Residues on the left side of the boxed region are corresponding to near the outer membrane clasp (OMC) of the α subunit and were negatively correlated with the beta transmembrane region in ITK and IT, while this correlation was eliminated in IK.

**Figure 3:**
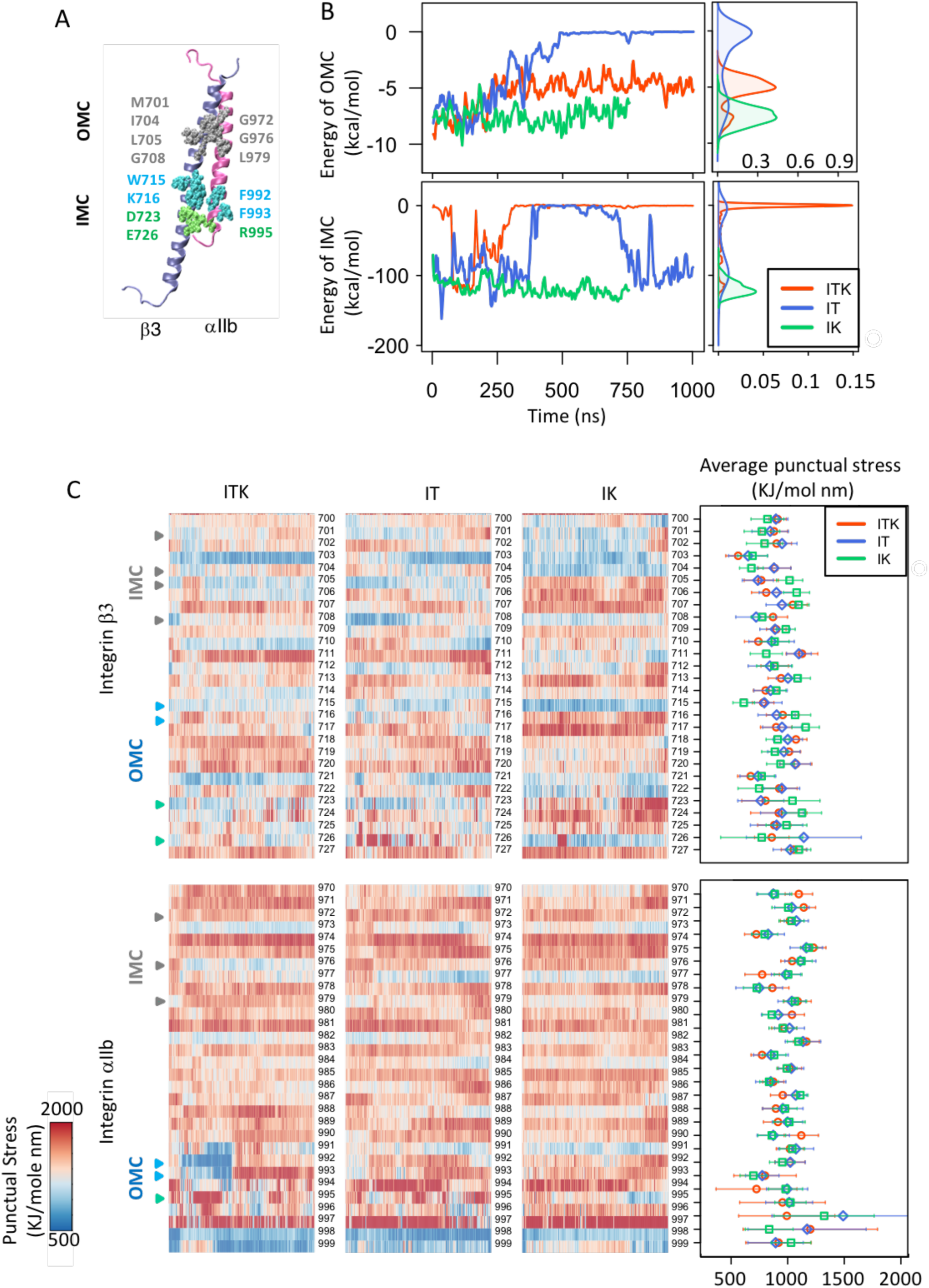
Interactions between αIIb and β 3 dimer. **(A)** Structure of the αIIb (mauve) β3 (iceblue) heterodimer. Interacting residues in the OMC region are shown in silver, in the IMC region in blue, and the αIIb R995 - β3 D723/D726 salt bridges in green. **(B)** The time plot and density plot of energy of interactions between OMC (top) and IMC (bottom) residues are shown for the ITK, IT, and IK simulations in red, blue, and green, respectively. **(C)** Time-resolved force distribution analysis (FDA) and time-averaged puntual stress within the integrin αIIb and β3 dimer were calculated. Representative heat maps of perresidue punctual stress values over simulation time for the integrin β3 residues 700 to 727 and αIIb residues 971 to 996 are shown. A representative heatmap is shown for each of the three ITK, IT, and IK simulations. Arrows highlight the location of IMC and OMC residues, which show significant changes in punctual stresses under force.

In addition, we calculated the crossing angle between αIIb and β3 helices (θ) as a function of time (Fig. 2D) to quantify the conformational changes of integrin. Specifically, we calculated the angle between the line crossing residues 967 to 979 in αIIb and the line crossing residues 697 to 709 in β3. As expected, an increase in crossing angle of β3 in IT simulations was observed. Interestingly, in ITK, the θ angle starts from 40 and then approaches zero after 250 ns and αIIb and β3 helices become almost parallel to each other. In the case of binding of kindlin2 without talin1 (IK), our results showed a small angle change (10 degrees) in θ, which is clearly in contrast with the cases of binding of talin1 that resulted in a noticeable change of θ (i.e., 30 degrees change in IT and 40 degrees change in ITK).

### Interactions between αIIb and β3 upon binding of talin1 and kindlin2

In order to determine the conformational transition in integrin β3 upon binding of talin1 and kindlin2, we next analyzed the interaction energy between αIIb and β3 helices. Two interaction interfaces on the integrin αIIbβ3 TM/CT domains are known to stabilize the resting conformation of the dimer: the inner membrane clasp (IMC) and the outer membrane clasp (OMC). IMC is characterized by the interactions between the highly conserved ^991^GFFKR^995^ motif of αIIb and W715, K716, and I719 residues of β3, and a salt bridge between residue R995 on αIIb and D723 on β3 (Fig. 3A). The R995-E726 salt bridge is also known to be involved in maintaining integrin in an inactive conformation. OMC is defined as an interaction network between ^972^GXXXG^976^ on αIIb and V700, M701, and I704 residues on β3, as well as associations between αIIb ^979^LL^980^ and β3 ^705^LXXG^708^ residues. Mutations in the residues involved in IMC or OMC result in disruption of interactions between αIIb and β3 helices leading to integrin activation (6,19,28,41,42).

To examine whether the binding of kindlin2 or talin1 could disrupt the IMC and OMC interactions in our simulations, we calculated the non-bonded interaction energy between IMC and OMC regions over the course of simulation time (Fig. 3B). Our results showed that the non-bonded interaction energies of IMC and OMC decrease upon simultaneous binding of talin and kindlin (ITK simulation) and stabilizes after 250ns approximately zero in IMC and −4 kcal/mol in OMC. In IT simulation, the nonbonded interaction energy in OMC approaches zero and remains zero after 500ns. On the other hand, IMC interactions in IT simulations switch between zero and −100kcal/mol. In IK simulation, the level of OMC and IMC both remain stable at −7 kcal/mol, and −120 kcal/mol, respectively. These results indicate that kindlin2 and talin1 cooperation can destabilize integrin αIIbβ3 dimer at IMC region and weaken their association in OMC region. Our results also suggest that kindlin2 alone is not sufficient to disrupt the IMC or OMC interactions.

Mechanical signal transmission entails dynamic force redistribution across the integrin αIIbβ3 molecule. To investigate how cytoplasmic interaction(s) with talin1 and/or kindlin2 allosterically impact the stress distribution within the transmembrane domain, especially near the IMC and OMC, we performed time-resolved force distribution analysis (FDA) as shown in Fig. 3D. The time-average of punctual stress of all integrin residues was used as a measure to compare force distribution patterns between simulations as shown in Fig. 3D, bars indicate standard deviation. Significant differences in the time-averaged punctual stress (>80 KJ/mol nm) were almost exclusively observed in IK compared to ITK and IT. Specifically, time-averaged punctual stresses experienced in residues 704 and 705 (part of IMC), 706, 711, 715 (part of OMC), 717, 722, 723 and 724 were remarkably higher in IK, while they were almost the same in IT and ITK. One exception was residue 726 for which time-averaged punctual stress in IT was higher than that in the ITK and IK simulations. Notably, residue 711 showed a most significantly different punctual stress within the IK simulation.

### αIIb-β3 angle change (θ) is correlated with OMC

To understand whether changes in the angle (θ) between αIIb and β3 (Fig. 2D) was correlated with the interaction energies of IMC and OMC (Fig. 3B), we performed a cross correlation analysis using the cross correlation function for time series in R, as described in **Methods** (Fig. 4). This function computes the covariance between two time series up to a defined lag. The θ angle was not a significant predictor of IMC, most likely because IMC was either abruptly disrupted or remained stable throughout the simulation. This suggests that IMC disassociation is directly regulated by cytoplasmic interactions and not through angle change. In other words, interaction at the cytoplasmic region exerts tension that is transmitted by both bonded and non-bonded interactions across integrin residues towards the membrane, which eventually breaks IMC and allows angle change. Conversely, dynamics of OMC interaction was highly correlated with θ in ITK and IT. The reverse correlations of ITK and IT reflects opposite directions of angle change (Fig. 4). Since there is no significant angle change (θ) in the IK simulation, we see a minimal θ-OMC correlation compared to ITK and IT.

**Figure 4:**
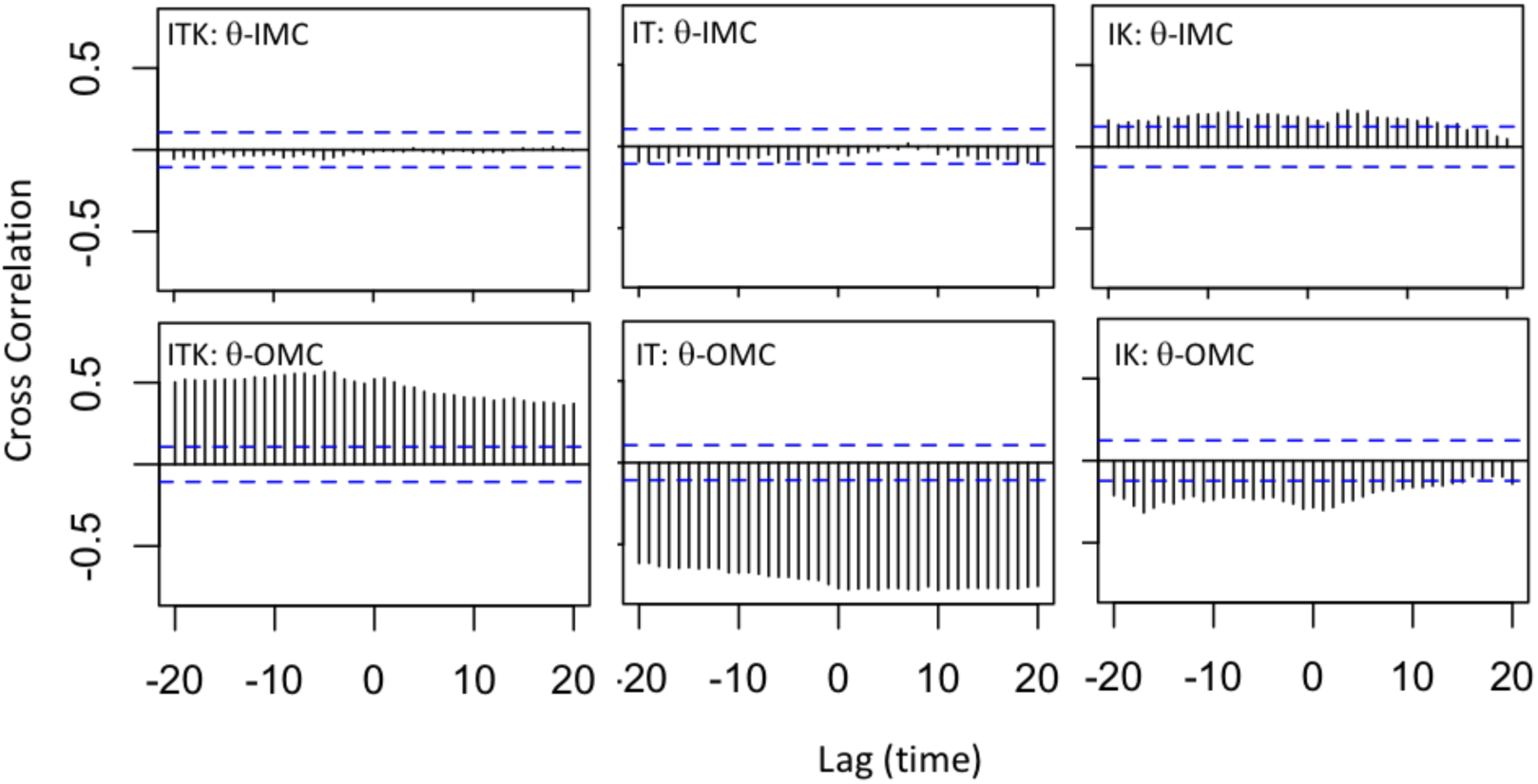
θ angle correlation with IMC and OMC. Cross correlation of IMC and angle changes of αIIb and β3 (top); and OMC and angle changes of αIIb and β3 (bottom) are shown. The horizontal blue lines are the approximate 95% confidence interval. Cross correlation plot is shown for each of the three ITK, IT, and IK simulations. High correlations exist between θ and OMC in ITK and IT, while the other correlations are negligible.

### Talin1 binds more effectively/strongly to integrin αIIbβ3 in the presence of kindlin2

Integrin β3 tail contains 47 residues and two NxxY motifs. The first NPLY motif (membrane proximal) is the binding site for talin1 and the second NITY motif (membrane distal) is the kindlin2 binding site. Talin1 and kindlin2 can simultaneously bind to β3 through their F3 domains and initiate the inside-out signaling (3). In order to understand how this inside-out signaling may be initiated by the cooperation between kindlin2 and talin1, we investigated the molecular mechanisms of simultaneous binding of talin1 and kindlin2 to the short CT of β3 and compared them with individual binding of integrin with talin1 and kindlin2 in our simulations. Specifically, we calculated non-bonded interaction energies between integrin β3 and kindlin2 or talin1, as well as solvent accessible surface areas of integrin β3 CT, talin1 and kindlin2.

We first calculated the non-bonded interaction energy between the CT of integrin β3 and talin1 or kindlin2 F3 domains (Fig. 5A). Specifically, we calculated the interaction between residues 720 to 762 of β3 and residues 569 to 680 of talin1 or residues 560 to 680 of kindlin2. In the IT simulations, talin and β3 maintain an average interaction energy of 105 kcal/mol (Fig. 5A top). However, talin1 is able to bind much more strongly to integrin β3 over time in ITK simulations. In this simulation, the nonbonded interaction energy between talin1 and β3 increases from 100 to 400 kcal/mol with an average energy of 222±94 kcal/mol. Kindlin2, on the other hand (Fig. 5A bottom) binds more strongly to integrin β3 in IK simulation with an interaction energy of 258±61 kcal/mol, compared to 124±57 kcal/mol in ITK simulation.

**Figure 5:**
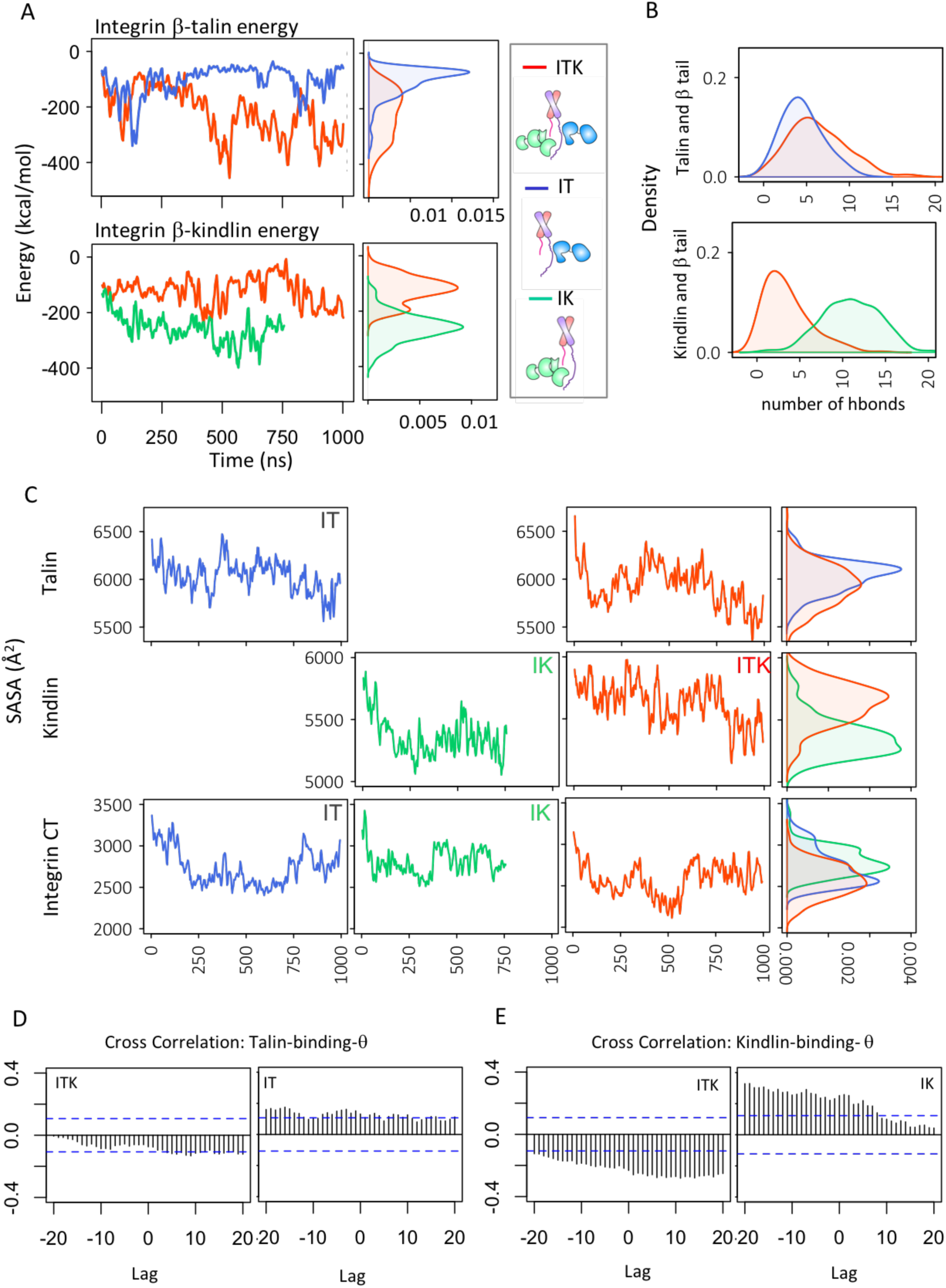
Integrin β 3-talin1 or kindlin2 binding. **(A)** The time and density plots of interaction energies between integrin β3 CT and talin1 F3 domain (top) and integrin β3 CT and kindlin2 F3 domain (bottom) are shown in three simulations of ITK, IT, and IK. **(B)** Density plot representations of the number of hbonds between integrin β3 CT and talin1 F3 domain (top) is shown for each of the ITK and IT simulations. The same plot is shown for kindlin2 (bottom) for each of the ITK and IK simulations. **(C)** Solvent accessible surface area (SASA) of talin1 (subdomain that binds integrin), in IT or ITK simulations (top). SASA of kindlin2 (subdomain that binds integrin), in IK or ITK simulations (middle). SASA of the CTs of integrin in IT, IK, or ITK2 simulations (bottom). **(D)** Cross correlation function between binding energy of talin1-integrin β3 and θ in ITK and IT simulations are shown. **(E)** Cross correlation between binding energy of kindlin2 – integrin β3 and θ in ITK and IK are shown (bottom).

Next, we calculated the solvent accessible surface area (SASA) of the integrin-binding-F3 subdomains of talin1 and kindlin2, and integrin β3 CT (see **Materials and Methods**, Fig. 5B). Our results show that the SASA of talin1 is lower in ITK simulations (5929±210 Å^2^) compared with IT simulation (6034±163 Å^2^) (Fig. 5A top). The SASA of kindlin2 immediately drops in the first 250ns of simulation in IK simulations. However, when talin1 is present, the SASA of kindlin2 fluctuates much more slowly and is much higher in value as shown in the density plots (Fig. 5B middle). The SASA of integrin CT is the lowest in ITK simulations compared to IT or IK simulations. The average values of 2575±189 for ITK, 2698±221 for IT, and 2805±158 for IK suggests that both can bind to various regions on integrin and reduce its SASA (Fig. 5B bottom). Also, in IT simulation, the SASA of integrin peaks at a slightly lower value than IK simulation, indicating a closer contact between integrin and talin1, compared to integrin and kindlin2 in conditions where only one of them is present. Based on the interaction energies and SASA results, it is evident that talin1 binds more effectively in the presence of kindlin2, and kindlin2 can bind to integrin much more strongly in the absence of talin1.

The crystal structures of the kindlin2-integrinβ3 and talin1-integrinβ3 complexes revealed the hbond network between the molecules (33). To determine the stability of kindlin2 or talin1-integrin complex, we calculated the number of hbonds between integrin CT - kindlin F3 and integrin CT - talin F3. Our results show that talin1 forms a slightly higher number of hbonds with integrin β3 tail in the presence of kindlin2. However, kindlin2 can form a larger number of hbonds with integrin β3 tail in the absence of talin1.

Finally, to determine whether the interaction energies between integrin αIIbβ3 CT and talin1 or kindlin2 were directly correlated with the changes observed in the crossing angle between αIIb-β3 (θ), we also calculated the cross correlation between these parameters. The interaction between talin1 and integrin β3 was promptly formed in both IT and ITK, while kindlin2 binding was relatively gradual. Kindlin2 binding lagged the angle change in IK, while in ITK, it did not strongly correlate with θ change. In both IT and ITK, talin binding did not show a major correlation with the angle change mainly because talin1 interaction was stable throughout the simulations. This suggests that a stable binding with talin1 is necessary for the subsequent changes in the integrin conformation. However, the role of kindlin2 is more complex as it may both directly and indirectly modulate integrin activation.

### Force distribution analysis of integrin β3 CT

We performed time-resolved force distribution analysis to monitor the dynamics of stress redistribution within the β3 cytoplasmic region upon binding to talin1 and kindlin2. We have calculated the punctual stresses, scalar pairwise forces exerted on each atom, for the residues 728 to 762 of integrin β3 cytoplasmic domain over time for both IT and ITK simulations (Fig. 6A). It can be seen from the heat map that the punctual stress fluctuates in most of the residues, except in residues 729 and 736, where the punctual stress remains high all the time both in IT and ITK.

**Figure 6:**
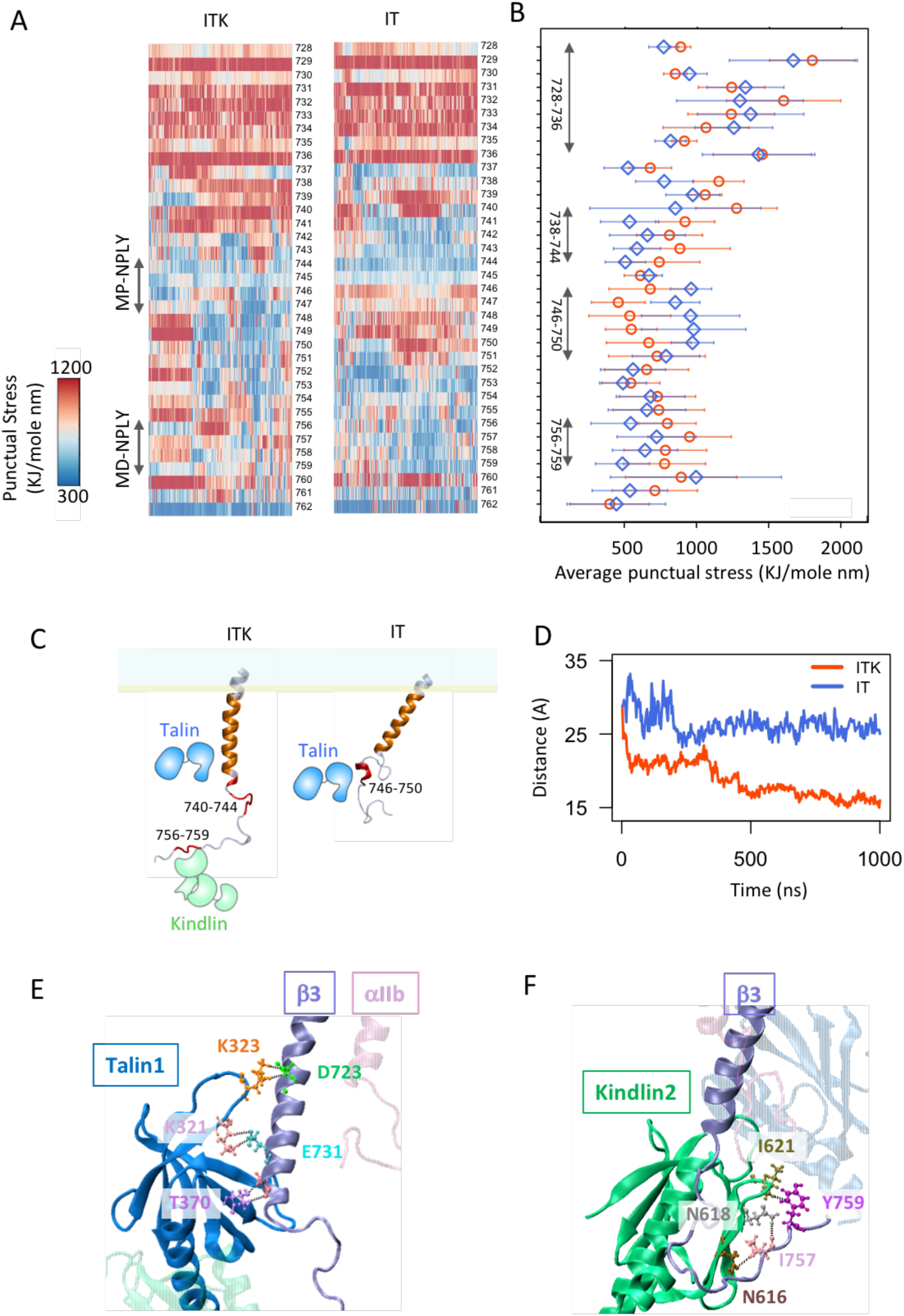
Force distribution analysis of integrin αIIbβ 3 CT. **(A)** TRFDA of integrin αIIb and β3 subunits. Representative heat maps of per-residue punctual stress values over simulation time for the integrin β3 cytoplasmic regions are shown. A representative plot is shown for each of the three ITK, IT, and IK simulations. **(B)** Schematic representations of the final structure of the integrin β3 CT in ITK and IT simulations. Regions with high punctual stress values are shown in red and orange on β3 CT. **(C)** The time plot of distance between the center of mass of residues 720 to 736 in integrin β3 and talin1 F3 subdomain are shown for the ITK and IT simulations in red, blue, respectively. **(D)** The time plot of distance between the center of mass of residues 720 to 736 in integrin β3 and talin1 F3 subdomain are shown for the ITK and IT simulations in red, blue, respectively. **(E)** Strong interactions between talin1 F3 subdomain and integrin β3 membrane proximal region in ITK simulation, mainly between residues K321, K323, and T370 on talin1, and residues D723, E731, and A735 on β3. Hydrogen bonds are shown with dashed black lines. **(F)** Interactions between kindlin2 F3 subdomain and integrin β3 NITY motif in ITK simulation, mainly between residues N616, N618, and TI621 on kindlin2, and residues I757 and Y759 on β3. Hydrogen bonds are shown with dashed black lines.

Also, we have calculated the time-averaged punctual stress over the last 300 ns of the simulations (Fig. B). Different distinct regions can be identified on Fig. 6B and 6C based on whether the punctual stress of IT or ITK dominates. Time-averaged punctual stress is higher in residues 746 to 750, which overlaps with the talin1 binding site (NPLY), in IT compared to ITK. Conversely, the kindlin2 binding site (NITY) consisting of residues 755 to 759 shows higher time-averaged punctual stress values in ITK compared to IT. This most likely indicates that direct binding with kindlin can locally increase stress levels. However, the region consisting of residues 738 to 744 exhibits higher average stress values in ITK compared to IT. We should note that in ITK, the stress value suddenly declines in residues 748 and 749, while at the same time it increases in residues 737 and 739.

Moreover, we see similar high stress values in the region consisting of residues 728 to 736. Our analysis of binding interactions in this region revealed that these residues interact strongly with the membrane in both IT and ITK. On the other hand, the distance between the center of mass of talin F3 subdomain and integrin β3 residues 728 to 736 decreases more significantly in ITK compared to IT (Fig. 6D). Therefore, the presence of kindlin resulted in significantly stronger interaction of talin with integrin β3 MP region in ITK simulation. This is most probably an important factor in integrin activation as can disrupt IMC.

Furthermore, we studied the interactions of talin1 and kindlin2 with high-stress regions of integrin β3 over the course of the ITK simulation. We observed three important interactions between the F3 subdomain of talin1 and the MP region of integrin β3 (residues 723 to 736), namely, 1) talin K323-β3 D723, 2) talin K321-β3 E731, and 3) talin T370-β3 A735 (Fig. 6E). In addition, interactions formed between the F3 subdomain of kindlin2 and the NITY motif of integrin β3 (residues 756 to 759). Specifically, residues N616/N618 and I621 of kindlin2 interacted with I757 and Y759 of integrin β3, respectively, which remained stable for the last 50ns of the ITK simulation (Fig. 6F).

### Distinct conformational changes of kindlin2 and talin1 in presence of each other

To determine whether any conformational changes were induced in kindlin2 or talin1 by their mutual interactions with integrin αIIbβ3, we performed structural alignments between the final structures of kindlin2 (at t=760ns) and talin1 (at t=1000ns) in each simulation, and their initial crystal structures. The structural alignments were performed using all sub-domains that were simulated (i.e. F2-F3 for talin1 and F0-F3 for kindlin2). However, for clarity, the alignments are shown separately for each sub-domain (Fig. 7). In order to quantify the deviation of each residue from its original position, the distance between residues of the initial and final structures were calculated after structural alignment was performed (Fig. 7). Our results show that the conformational changes observed in the F2 subdomain (residues 196 to 305) of talin1 are much higher in IT simulation compared with ITK simulation (Figure 7A). These conformational changes are mainly observed in two α helices (∼ residues 224 to 294) in the four helix bundle of the F2 subdomain in IT simulation. The conformational changes observed in the F3 subdomain of talin1 were similar between IT and ITK conditions and mostly limited to the loop regions as expected. The structural alignment of kindlin2 showed significant conformational changes in the F0 subdomain in IK simulations, and F3 in ITK simulations, whereas the F1 and F2 subdomains showed very small changes in both simulations. Previous work showed that the F0 subdomain of kindlin2 is highly dynamic and moves independently of F1-F3 subdomains which could be a reason for the difference in the conformational state of F0 between different simulations. On the other hand, the conformational changes in F3 were seen in integrin binding regions (i.e. residues 590 to 656) and are likely attributed to the distinct interactions of kindlin2 with integrin β3 in IK vs ITK simulations. These results suggest that conformational changes are induced in talin1 and kindlin2 upon interactions with integrin β3, and that these changes are distinct when both molecules are present.

**Figure 7:**
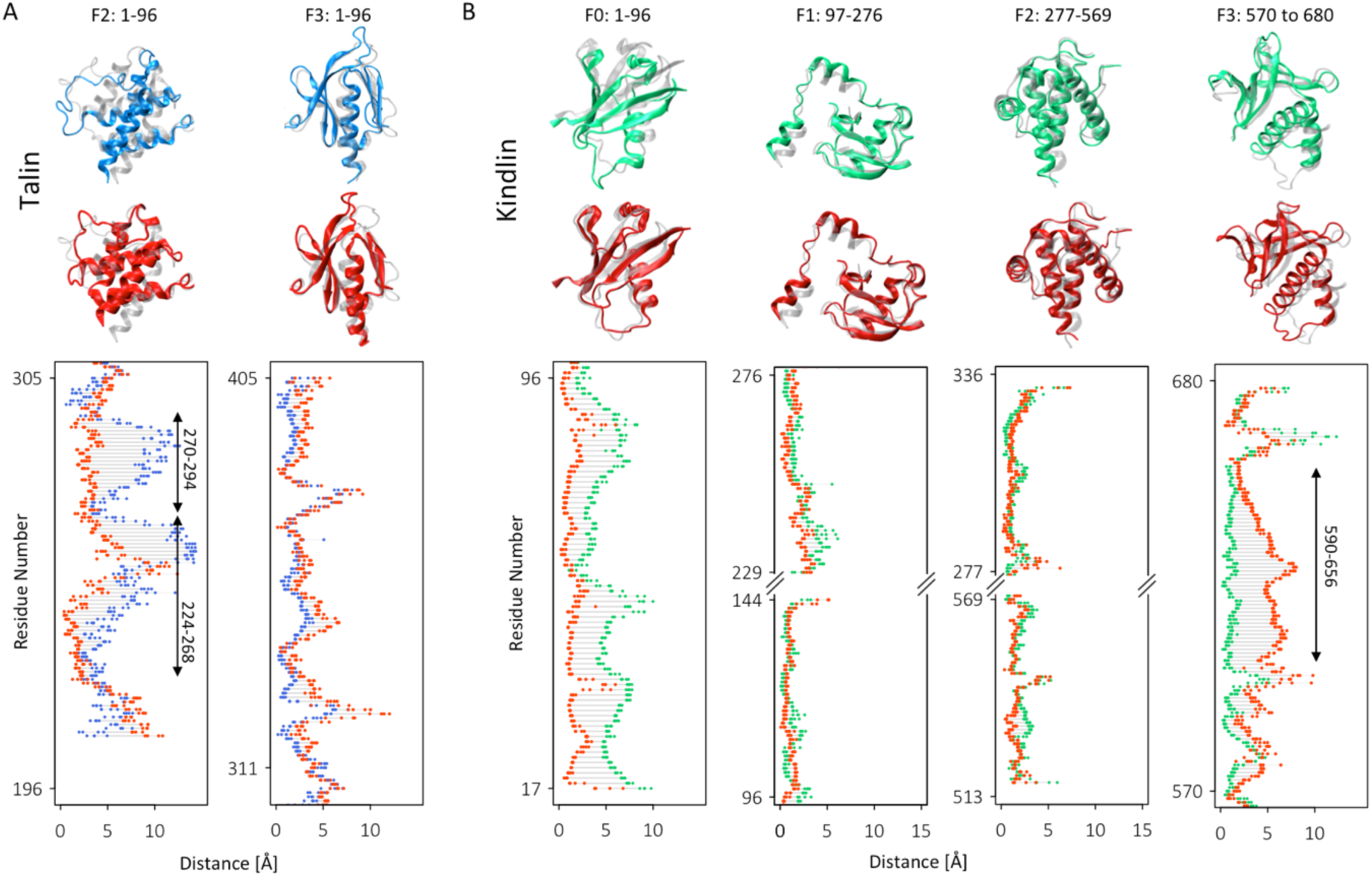
Kindlin2 and talin1 structural alignment. **(A)** Final frames of talin1 in IT (blue) and ITK (red) were aligned to initial pdb of talin1 (transparent). The distance shows the 3D distance between each residue in the two structures after alignment. **(B)** Final frames of kindlin2 in IK (green) and ITK (red) were aligned to the initial pdb of kindlin2 (transparent) (by RMSD minimization methods). The distance shows the 3D distance between each residue in the two structures after structural alignment.

## Discussion

Integrin activation is central to a variety of cell functions, including cell-matrix adhesion, cell migration, and apoptosis. Kindlin and talin are both cytoplasmic proteins that directly interact with integrin tail and induce a high-affinity state of integrin for extracellular ligands (3). Although talin can induce conformational changes in integrin, it has been shown that talin alone is not sufficient for the activation of integrin and kindlin also plays a key role in this process (14,17). While molecular mechanisms of talin-mediated integrin activation have been extensively studied, it is not yet clear how kindlin interferes with this process. Kindlin dysfunction has been associated with a number of human diseases, which are likely linked to deficiency in integrin activation; thus knowing the molecular mechanism of kindlin-mediated integrin activation has potential impacts in therapeutic purposes. Here, we investigated the interplay between talin and kindlin in integrin activation using microsecond long molecular dynamics simulations (Fig. 1).

Previous studies have shown that kindlin binding to integrin is not sufficient but is required for triggering integrin activation (6,13,15). However, the mechanisms of kindlin’s involvement remain elusive, in particular it is still not known whether kindlin can induce conformational changes to the integrin heterodimer. To address these questions and provide molecular details of the interaction among kindlin and talin with integrin, we developed all-atom microsecond-scale MD simulations of αllbβ3 using an explicit lipid-water environment under three distinct scenarios, namely integrin in complex with talin1 (IT), kindlin2 (IK), and both talin and kindlin (ITK). Our control simulations featuring integrin αIIbβ3 and kindlin2 (IK) showed that the main interactions that keep integrin in an inactive state, namely the IMC and OMC, remained stable over the course of 750 ns of the simulation when only kindlin is present (Fig. 3B). This further confirmed that kindlin binding is insufficient for unclasping cytoplasm and transmembrane regions of αIIbβ3.

Interestingly, our simulations showed that kindlin2 binds to the integrin β3 tail more strongly in the absence of talin1, whereas talin1 binding to integrin is reinforced by kindlin2 (Fig. 5). Talin1 interaction was particularly enhanced with the membrane proximal region of integrin β3 (residues 720 to 736), while its interaction with the membrane MP NPXY motif was gradually weakened over time. Therefore, kindlin2 physically pushes talin1 towards the membrane resulting in a more effective disruption of the IMC. This way, kindlin2 modifies the activation pathway by positioning talin1 more upward and closer to the membrane, resulting in stronger binding and a complete IMC disruption. Also, kindlin2 decreased the intradomain conformational changes of talin1, especially in F2 subdomain, which promoted both talin1-lipid and talin1-integrin β3 CT interactions (Fig. 7A). Therefore, we suggest that the strength of kindlin2 binding to integrin does not necessarily correlate with activation, while stronger talin1 association can expedite the activation process. This further confirms the indirect role of kindlin2 as a cooperative partner for talin1 rather than a direct activator of integrin αIIbβ3. It should be noted that even though we did not observe a strong interaction between talin1 and membrane proximal region of β3 in IT simulations, it is quite conceivable that even without kindlin2’s assistance, talin1 can eventually bind to the proximal region of integrin tail in a longer timescale. On the other hand, the stronger binding of kindlin2 to integrin in the absence of talin1 observed in our simulations may have implications in talin-independent roles of kindlin in the formation and maturation of focal adhesions. Theodosiou et al. showed that subsequent to the cooperative activation of integrin by talin and kindlin2, kindlin2 initiates a talin-independent signaling pathway at new adhesion sites by binding to other molecules such as paxillin to induce cell spreading (15). Based on previous studies and our results, we propose that kindlin cooperates with talin to activate integrins by enhancing talin’s interactions with the CT of integrin. On the other hand, kindlin can bind more strongly to integrin in the absence of talin to perform talin-independent signaling subsequent to integrin activation (15).

It is now widely accepted that the inner membrane clasp (IMC) and outer membrane clasp (OMC) between integrin subunits maintains the resting state of integrin and unclasping it triggers the inside-out signaling (6). In the IMC, R995 of beta-integrin forms two salt bridges with D723 and E726 on alpha-integrin, however, it has been shown that disrupting the R995-D723 interaction is most critical for integrin activation (6). Our simulations indicated that talin1 disrupts R995-D723 interaction in both presence or absence of kindlin2 (Fig. S1), however, kindlin2 is also required for dissociating R995-E726 within 1μs of simulation (Fig. 3B). Conversely, the OMC was destabilized in both cases (IT and ITK), causing significant conformational changes in the transmembrane region of integrin αIIbβ3 (Fig. 2 and 3). The destabilization of OMC is highly correlated with the dramatic angle change between αIIb and β3 (θ) in both IT and ITK (Fig. 4). We noticed that the angle change in ITK was in reverse direction relative to that in IT, i.e., decreasing angle in ITK and increasing angle in IT (see Fig. 2). Previous studies suggested that talin-mediated integrin activation involved an increase in the αIIb-β3 crossing angle and subsequent separation of the transmembrane domains (7,27). However, recent studies showed that transmembrane domains of integrin subunits were close to each other in all conformational snapshots including active, inactive and intermediate states, contradicting the notion of chain separation as an indicator of integrin activation (28,43). Since the angle change in ITK led to disruption of the IMC and OMC interactions, we propose that the degree of angle change is more pivotal in integrin activation than the direction of such change. This is most likely because talin1 and kindlin2 eliminate the inhibitory effects of IMC and OMC, which allows for mechanical signals to be transmitted across the membrane, without complete separation of the transmembrane regions. This is also demonstrated by changes in the force distribution patterns of the transmembrane domains of integrin in both IT and ITK simulations (Fig. 3). Furthermore, we showed previously that residue A711 in β3 helix is key for signal transmission across the transmembrane domain of integrin and an A711P mutation can disrupt integrin activation (44). Our FDA analysis also indicated that A711 bears high forces in both IT and ITK but not in IK (Figure 3), implying that signal transmission is activated upon IMC disruption. Overall, our study suggests that not only both talin1 and kindlin2 are essential for efficiently initiating the activation process, but also kindlin2 may modify the molecular mechanism of inside-out signaling at the integrin aIIbb3 tail, which eventually leads to the opening of the integrin ectodomain.

Although not studied in this work, it is important to note that kindlin may have several other roles in integrin activation and the formation of focal adhesions. For example, one possible role of kindlin in integrin activation may be preventing the association of cytoplasmic inhibitors of integrin activation such as filamin or α-actinin to the integrin tail. Also, kindlin may be involved in integrin clustering through dimerization (31). Moreover, kindlin binding to actin can enhance cytoskeletal coupling to the membrane. Therefore, simultaneous binding of kindlin and talin to the integrin β increases the overall force applied to the integrin tail, which may facilitate focal adhesion formation and maturation.

Taken together, our results shed light on the molecular mechanisms by which kindlin2 cooperates with talin1 in integrin αIIbβ3 activation. This is the first computational study, to the best of our knowledge, that models the interplay between kindlin2 with talin1 in promoting inside-out signaling through integrin binding.

## Author Contributions

All authors contributed to the design and implementation of the project and the computational experiments. All authors provided critical feedback in all steps of the project, which shaped the final research. ZH conducted the molecular dynamics simulations with inputs from HS and ZJ. All authors were involved in post-processing and analysis of the data and writing of the manuscript. MRKM supervised the project and contributed materials and analysis tools.

### Acknowledgments

We acknowledge fruitful conversations with Mehrdad Mehrbod in the early stages of this research. This research used resources of the National Energy Research Scientific Computing (NERSC) Center, a Department of Energy Office of Science user facility supported by the Office of Science of the United States Department of Energy under contract no. DE-AC02-05CH11231.

